# *HOXB7* induces STAT3-mediated transformation and lung metastasis in immortalized mammary gland NMuMG cells

**DOI:** 10.1101/2021.05.16.444388

**Authors:** Kazushi Azuma, Mai Sakamoto, Shota Katayama, Atsuka Matsui, Kazuya Nakamichi, Naoki Goshima, Shinya Watanabe, Jun Nakayama, Kentaro Semba

**Author notes:** Corresponding authors **Correspondence to:** Jun Nakayama (or) and Kentaro Semba, Department of Life Science and Medical Bioscience, School of Advanced Science and Engineering, Waseda University, TWIns, 2-2 Wakamatsu-cho, Shinjuku-ku, Tokyo, 162-8480, Japan. These authors equality contributed (Kazushi Azuma, Mai Sakamoto, Shota Katayama).

## Abstract

The homeobox family genes are often dysregulated in a various cancer type. Particularly *HOXB7* amplification and overexpression correlate with poor prognosis in various cancer such as gastric, pancreatic, and lung cancers. Moreover, *HOXB7* is known to contribute to cancer progression by promoting epithelial to mesenchymal transition, anti-cancer drug resistance, and angiogenesis. In this study, we show that *HOXB7* is coamplified with *ERBB2* in a subset of breast cancer patients and HOXB7 expression correlates with poor prognosis in HER2-positive breast cancer patients. This clinical observation is supported by the following results: *HOXB7* overexpression in an immortalized murine mammary gland epithelial cell line NMuMG induces cellular transformation *in vitro*, tumorigenesis and lung metastasis through the activation of JAK-STAT signaling.

## 1. Introduction

Oncogenic alterations, such as gene amplifications, mutations, chromosomal translocation, and epigenetic dysregulation, cause tumorigenesis *in vivo* (Hanahan & Weinberg 2011). Activated oncogenes also induce typical phenotypic changes, so-called transformation, in immortalized normal cells such as a NIH3T3 and normal murine mammary gland (NMuMG) cells *in vitro*. Therefore, assay systems evaluating such phenotypic changes, namely focus formation, soft agar colonization, and tumorigenesis have been used for the screening of novel oncogenes in previous studies including ours (Saito *et al*. 2012; Matsui *et al*. 2016).

Most oncogenes cause dysregulation of the signal transduction cascades that relate to proliferation, cell cycle, and survival (Kolch *et al*. 2015). Typically, oncogenic signaling pathways, such as RAS-MAPK and PI3K-AKT are often activated in various cancer cell types and thus molecular targeted-drugs against them have been successfully developed. Recent studies showed that the JAK-STAT signaling is also important in cancer initiation and progression (Hu *et al*. 2021). STAT3 is activated in cancer stem cell-like populations (Marotta *et al*. 2011), and regulates stemness via fatty metabolism in breast cancer (Wang *et al*. 2018). In addition, aberrant activation of STAT3 with IL6-enriched tumor microenvironment is associated with an immunosuppression of tumor-infiltrating immune cells (Qureshy *et al*. 2020). JAK-STAT signaling also coactivates and supports cancer growth and progression with the activation of RAS-MAPK and PI3K-AKT pathways (Fazilaty & Mehdipour 2014).

The homeobox (HOX) genes encode transcriptional factors, that controls development and organ homeostasis. The HOX superfamily of homeobox genes (HOXA, HOXB, HOXC and HOXD clusters) are often dysregulated in breast, prostate, colon, and lung cancer. The differential dysregulation of HOX family genes in various solid tumors provides an opportunity to understand cancer development (Bhatlekar *et al*. 2014; Li *et al*. 2019). The HOXB clusters is composed of *HOXB1, HOXB2, HOXB3, HOXB4, HOXB6, HOXB7, HOXB8, HOXB9*, and *HOXB13* at 17q23.32. HOXB cluster genes are co-amplified by gene amplification in several types of cancer. Notably, overexpression of *HOXB7* contributes to cancer progression in breast, hepatocellular, oral, and gastric cancer cell lines (Errico *et al*. 2016; He *et al*. 2017; Wang *et al*. 2017). *HOXB7* induces the activation of TGFß signaling, epithelial to mesenchymal transition (EMT) and migration in breast cancer (Wu *et al*. 2006; Liu *et al*. 2015). In addition, the luminal subtype breast cancer cell line MCF7 acquires tamoxifen resistance via *HOXB7* expression (Jin *et al*. 2012). *HOXB7* as a cofactor of estrogen receptor (ER) causes cancer progression through the expression of downstream genes (Jin *et al*. 2015). In this study, we revealed HOXB7-mediated enhancement of JAK-STAT pathway and its possible involvement in cellular transformation *in vitro*. We also found that HOXB7 is enough to induce tumorigenesis and metastasis *in vivo* in the context of immortalized NMuMG cells.

## 2. Materials and methods

### 2.1. Cell culture

Normal murine mammary gland (NMuMG) and NMuMG-luc cells were cultured in Dulbecco’s modified Eagle’s medium (DMEM; Fujifilm Wako, Osaka Japan) supplemented with 10% fetal bovine serum (FBS; NICHIREI BIOSCIENCES INC., Tokyo, Japan), 100 U/mL penicillin G (Meiji-Seika Pharma Co., Ltd., Tokyo, Japan), 100 μg/mL streptomycin (Meiji-Seika Pharma Co., Ltd.), 10 μg/mL insulin (Fujifilm Wako), and 0.45% glucose (Fujifilm Wako) at 37°C with 5% CO_2_. Plat-E cells for retroviral packaging were cultured in DMEM supplemented with 10% FBS, 100 U/mL penicillin G, and 100 μg/mL streptomycin at 37°C with 5% CO_2_. Cells were treated with 0.1-1 μM ruxolitinib (LC Laboratories, MA, USA) under experimental conditions.

### 2.2. Retroviral packaging and infection

To establish NMuMG-Venus, *HOXB7*-expressing NMuMG (NMuMG-HOXB7), *HOXB7*-expressing NMuMG-luc (NMuMG-luc-HOXB7) cell lines by using retroviral infection, Plat-E cells were seeded into 6-well plates and transfected with pMXs-Venus-IRES-Puromycin^R^ and pMXs-*HOXB7*-IRES-Puromycin^R^ plasmid DNAs using 6 μg polyethylenimine (PEI; Polysciences, Inc., PA, USA). Then, the cell supernatants and 800 μg/mL polybrene (SIGMA ALDRICH Co., MO, USA), as previously described (Matsui *et al*. 2016). To select the expressing cells, 1 μg/mL puromycin (Fujifilm Wako) was added to the culture medium for 2 days.

To establish NMuMG-luc-HOXB7-shGFP, NMuMG-luc-HOXB7-shSTAT3#2 and NMuMG-luc-HOXB7-shSTAT3#3 cell lines by using retroviral infection, pSuper-shGFP-Hygromycin^R^, pSuper-shSTAT3#2-Hygromycin^R^ and pSuper-shSTAT3#3-Hygromycin^R^ plasmid DNAs were used with the method described above. To select the expressing cells, 800 μg/mL hygromycin was added to the culture medium for 5 days. The following pairs of oligonucleotides were used to target specified genes: shGFP, ACAACAGCCACAACGTCTATAGTGTGCTGTCCTATAGACGTTGTGGCTGTTGTTTTTTT and AAAAAAACAACAGCCACAACGTCTATAGGACAGCACACTATAGACGTTGTGGCTGTTGT; shSTAT3#2, CCTGGGTTGAATTGTCAGTTTGTGTGCTGTCCAAGCTGATAATTCAACTCAGGTTTT TGGAA and TTCCAAAAACCTGAGTTGAATTATCAGCTTGGACAGCACACAAACTGACAATTCAAC CCAGG; shSTAT3#3, CGACTTTGGTTTCAATTACGAGTGTGCTGTCCTTGTAGTTGAAATCAAAGTCGTTTTTGGAA and TTCCAAAAACGACTTTGATTTCAACTACAAGGACAGCACACTCGTAATTGAAACCAAAGTCG.

### 2.3. Western blotting

The infected cell lines were lysed with SDS sampling buffer (3.5 % SDS, 12% glycerol, 0.05M Tris-HCl (pH 6.8), 0.003% bromo phenol blue, and 5% 2-mercaptoethanol) for western blotting. Western blotting was performed as previously described (Matsui *et al*. 2016) with the antibodies shown in Supplementary Table S1. The cells were treated with 1μM ruxolitinib for 24 hours before harvesting.

### 2.4. Cell proliferation assay

A total of 2.5 × 10^4^ of NMuMG-Venus cells or NMuMG-HOXB7 cells were seeded into a 12-well plate. Cells were harvested with 0.25% trypsin (Thermo Fisher Scientific, MA, USA) and counted at each time point (24, 48 and 72 h) (Kuroiwa *et al*. 2020). Cells were continuously treated with ruxolitinib from 24 hours after cell seeding.

### 2.5. Focus formation assay

For focus formation assay, 7.0 × 10^4^ parental NMuMG cells and 3.5 × 10^4^ NMuMG-HOXB7 cells were mixed and seeded into 12-well plates. In STAT3 knock down experiment, 1.5 × 10^5^ of parental NMuMG cells and 7.5 × 10^2^ of NMuMG-luc-HOXB7 cell lines (NMuMG-luc-HOXB7-shGFP cells or NMuMG-luc-HOXB7-shSTAT3#3 cells) were mixed and seeded into 12-well plates. Cells were stained with 0.01% crystal violet/70% methanol to detect the cellular foci (Matsui *et al*. 2016). Cells were continuously treated with ruxolitinib from 24 hours after cell seeding. Photos were taken at the indicated day described in each figure legend.

### 2.6. Tumorigenesis assay

A total of 1.0 × 10^6^ NMuMG-Venus cells and NMuMG-HOXB7 cells were transplanted into 6-8-weeks-old *Rag2*^-/-^ female mice with 100 μL 50% Matrigel (BD Bioscience, CA, USA) as previously described (Ihara *et al*. 2017). All animal experiments were approved by the Animal Committee of Waseda University (WD19-058, 2019-A068).

### 2.7. Orthotopic Transplantation and bioluminescence imaging

A total of 1.0×10^6^ NMuMG-luc-HOXB7 cells, NMuMG-luc-HOXB7-shGFP cells, or NMuMG-luc-HOXB7-shSTAT3#3 cells were transplanted into fourth fat pads of 6-weeks-old BALB/cSlc-nu/nu female mice as previously described (Nakayama *et al*. 2017; Nakayama *et al*. 2022). After the injection, the transplantation sites were palpated for every 2-3 days to measure tumor formation time and mice were monitored primary tumor formation and metastasis formation by *in vivo* imaging system (IVIS Lumina Xr3; Perkin-elmer, Waltham, MA, USA) for every week. Mice were intraperitonially injected with 200 μL D-Luciferin (15 mg/mL; Gold Biotechnology, Inc., St. Luis, MO, USA), and Luminescence was taken under anesthesia using IVIS. Sixteen weeks after injection, all mice were sacrificed, and excised lungs. Lungs were transferred to a 12-well plate with 900 μL of D-PBS (-) (Fujifilm Wako Pure Chemical Industries) and 100 μL of D-Luciferin (15 mg/mL), and then luminescence was taken by IVIS, as previously described (Han *et al*. 2020). All animal experiments were approved by the Animal Committee of Waseda University (WD21-082, 2021-A074)

### 2.8. Quantitative real-time PCR

Total RNA was extracted using ISOGEN (NIPPON GENE Co., LTD, Tokyo, Japan). cDNA synthesis was performed using the Superscript III first-strand synthesis system (Thermo Fisher Scientific). Quantitative RT-PCR was performed using THUNDER BIRD SYBR qPCR mix (TOYOBO, Osaka, Japan) and 18S rRNA was used as a relative control. Primers are listed in Supplementary Table S2.

### 2.9. Clinical database analysis

The genomic profiles and RNA-seq datasets of The Cancer Genome Atlas (TCGA) breast cancer (BRCA) cohorts were subjected to OncoPrint and expression analyses. OncoPrint of *HOXB7* and *ERBB2* was conducted using cBioPortal (Cerami *et al*. 2012). The Kaplan-Meier method was used perform survival analyses of the molecular taxonomy of breast cancer international consortium (METABRIC) (Pereira *et al*. 2016), GSE1456 and GSE9893 data sets, as previously described (Han *et al*. 2022).

### 2.10. Statistical analysis

Welch’s t-test, Fisher’s exact test and log-rank test were performed using R 3.4.1 (https://www.r-project.org/).

## 3. Results

### 3.1. Expression of *HOXB7* correlated to poor prognosis in the HER2-positive breast cancer patients

To elucidate the clinical significance of *HOXB7* in breast cancer, we first analyzed RNA-seq data of TCGA BRCA cohorts for *HOXB7* expression. Gene expression analysis in each breast cancer subtype revealed that the HER2-positive subtype significantly highly expressed *HOXB7* compared with the other subtypes and normal tissue (Figure 1a). To examine the genomic profiles of *HOXB7* in TCGA BRCA cohorts, the mutational status of *ERBB2* (the gene that encodes HER2) and *HOXB7* was illustrated as an OncoPrint by cBioPortal (Figure 1b). *HOXB7* was significantly co-amplified with *ERBB2* in 30 patients out of 48 patients with *HOXB7* amplification (Figure 1c). HOXB7 expression also significantly increased in the *ERBB2/HOXB7*-amplified patients compared with the *ERBB2*-amplified patients (Figure 1d). These results suggested that *HOXB7* was preferentially amplified and expressed in HER2-positive breast cancer patients.

**Figure 1.**
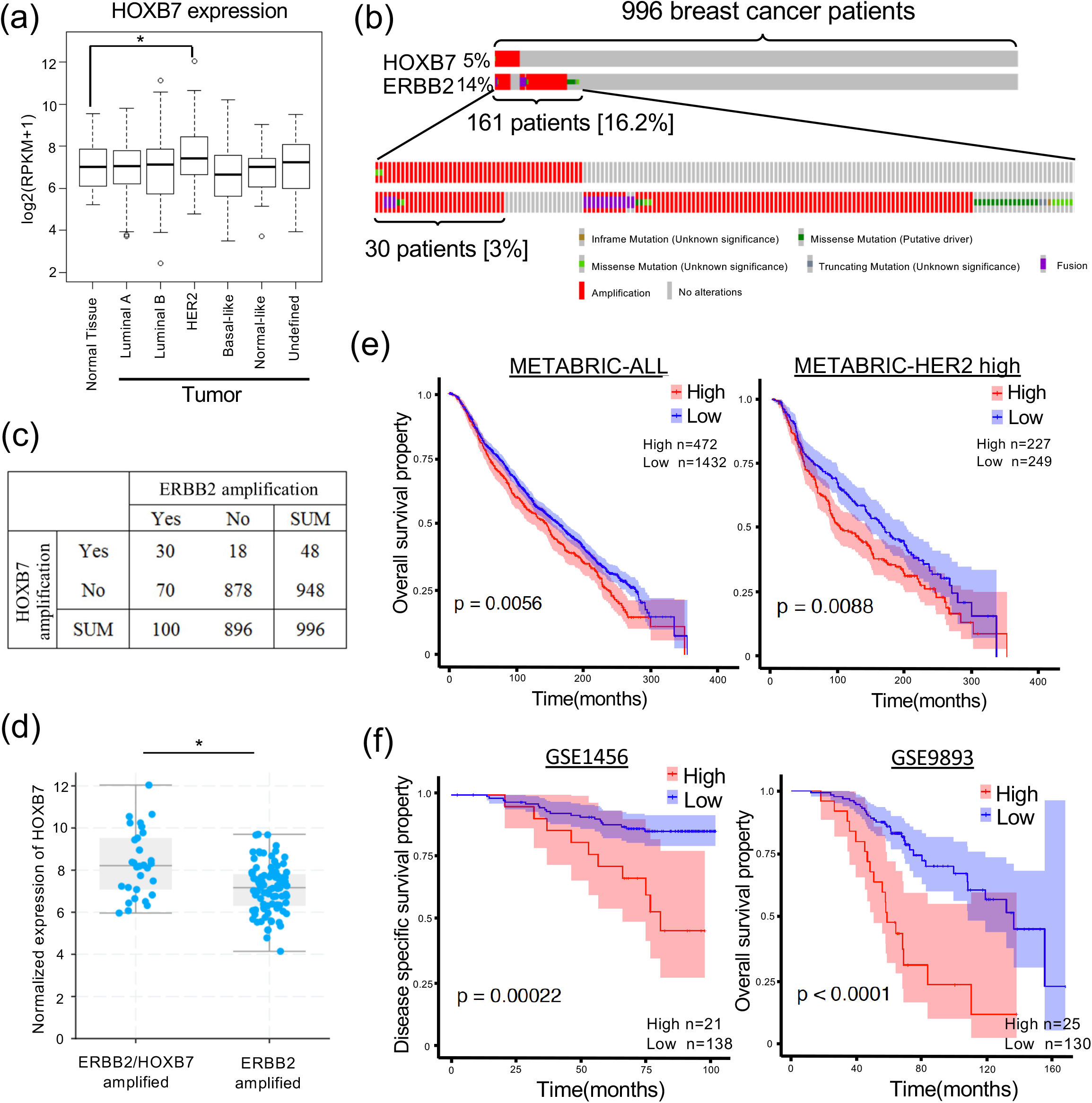
High expression of HOXB7 correlated to poor prognosis in breast cancer patients. (a) Comparative expression analysis of *HOXB7* in breast cancer subtypes of TCGA. Luminal A (n = 499), luminal B (n = 197), HER2-positive (n = 78), basal-like (n = 171), normal-like (n = 36), unidentified (n = 98). Welch’s t-test (p < 0.005). (b) OncoPrint of *ERBB2* and *HOXB7* genomic status in TCGA. (c) Altered genetic profiles of *ERBB2* and *HOXB7*. Fisher’s exact test (p < 0.05). (d) Comparative expression analysis of *HOXB7*. Welch’s t-test (p < 0.05). (e left) Survival analysis of all METABRIC cohorts with *HOXB7* expression and the overall survival status shown in a Kaplan-Meier plot. Shading along the curves shows 95% confidential interval. Log-rank test (p = 0.0056) (e right) METABRIC cohorts with high HER2 expression (476 patients; Top 25% of all METABRIC cohorts) Log-rank test (p = 0.0088) (f) Survival analysis of other breast cancer cohorts (GSE1456; disease specific survival; GSE9893; overall survival). GSE1456, log-rank test (p = 0.00022); GSE9893, log-rank test (p < 0.0001).

Next, to examine the relationship between *HOXB7* expression and progression in breast cancer, we performed a survival analysis using several public clinical data sets. In METABRIC cohorts, high expression of *HOXB7* was correlated with poor prognosis in all breast cancer patients (Figure 1e left) and especially in HER2-high patients (Figure 1e right). The correlation of poor prognosis and *HOXB7* expression was further confirmed by an analysis of other public breast cancer cohorts that included all subtypes (Figure 1f).

### 3.2. *HOXB7* induced oncogenic focus formation and activation of JAK-STAT3 signaling

To elucidate the oncogenic function of *HOXB7*, we first established NMuMG-HOXB7 cells that expressed *HOXB7* in an immortalized murine breast epithelial cell line NMuMG as a model system for both *in vitro* and *in vivo* evaluation of oncogenes (Matsui *et al*. 2016; Ihara *et al*. 2017). Expression of HOXB7 changed morphology of NMuMG cells from epithelial-like to spindle-like shape (Figure 2a). Furthermore NMuMG-HOXB7 cells formed foci (Figure 2b).

**Figure 2.**
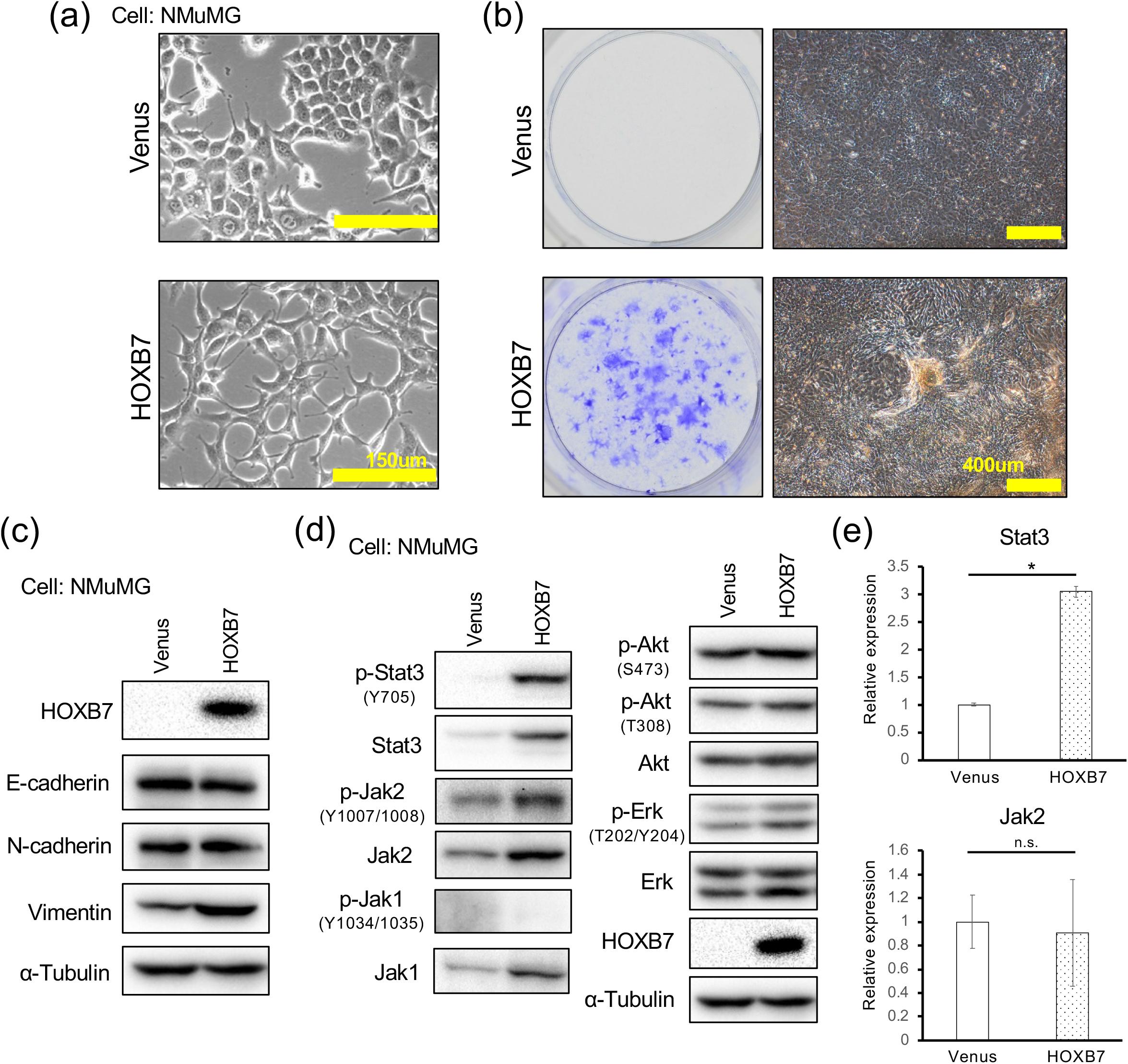
*HOXB7* expression induced transformation and activation of JAK-STAT signaling. (a) Bright imaging of *HOXB7*-expressing cells. (b) Focus formation assay in *HOXB7*-expressing NMuMG cells. *Venus*-expressing NMuMG cells were used as a negative control. Left: crystal violet staining. Right: bright field focus. Photos were taken at 10 days after the start of the experiment (n = 1 biological replicates; 3 independent experiments). (c) Western blotting analysis of EMT-related genes in *HOXB7*-expressing NMuMG cells. Venus-expressing NMuMG cells were used as a negative control. (d) Western blotting of JAK-STAT and growth signaling pathway in *HOXB7*-expressing cells. (e) qRT-PCR of *Stat3* and *Jak2*. Welch’s t-test (* p < 0.001, n = 3).

Next, to identify oncogenic signaling pathways of *HOXB7*, we performed western blotting using antibodies against major cancer-related signaling proteins. Among them, we detected the *HOXB7* overexpression increased the protein level of vimentin (a mesenchymal marker gene) (Figure 2c), but did not change those of E-cadherin and N-cadherin, suggesting *HOXB7* induces partial epithelial-mesenchymal transition. Notably, *HOXB7* overexpression drastically increased Stat3 phosphorylation and protein levels of Jak1, Jak2, and Stat3 (Figure 2d). Major proliferative signals, such as Erk and Akt, were not changed by *HOXB7* expression. Stat3 was increased at the transcriptional level, while Jak2 was increased at the protein level (Figure 2e).

Additionally, phosphorylation of Stat5 was also increased by *HOXB7* expression (Figure 3a). Phosphorylation of Stat3 and Stat5 induced by *HOXB7* expression was completely inhibited by a JAK1/2 inhibitor, ruxolitinib in a dose-dependent manner confirming JAK1/2 involvement in Stat3 and Stat5 phosphorylation (Figure 3a). Next, to elucidate the contribution of the oncogenic activity of p-Stat3, we performed *in vitro* proliferation assays and focus formation assays following ruxolitinib treatment. *HOXB7* decreased the proliferative activity of NMuMG cells *in vitro*, however, ruxolitinib treatment did not affect the proliferative activity of NMuMG-HOXB7 cells (Figure 3b). Nevertheless, ruxolitinib inhibited their focus-forming activity (Figure 3c). These results suggest that the activation of JAK-STAT signaling by *HOXB7* induced cellular transformation.

**Figure 3.**
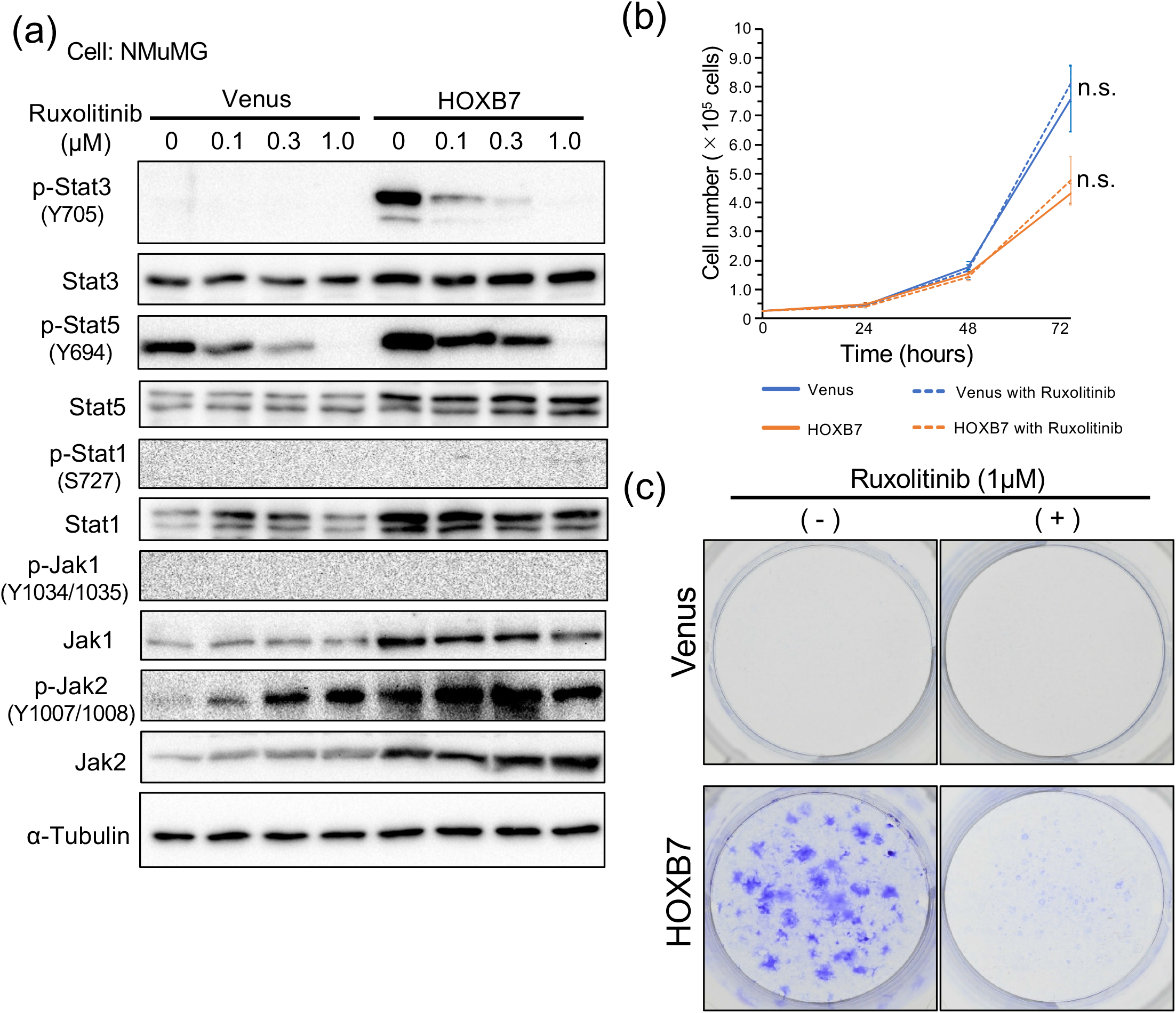
Focus formation by *HOXB7* was regulated by Stat3 activation. (a) Western blotting of JAK-STAT signal pathway in *HOXB7*-expressing NMuMG cells with ruxolitinib treatment. (b) *In vitro* growth curve of *HOXB7*-expressing NMuMG cells treated with ruxolitinib. (n.s.: Not significant, n = 3 biological replicates) (c) Focus formation assay of *HOXB7*-expressing NMuMG cells treated with ruxolitinib. Foci were stained with crystal violet. Photos were taken at 10 days after the start of the experiment (n=3 biological replicates; 2 independent experiments).

### 3.3. Overexpression of *HOXB7* induced tumorigenesis and metastasis

To examine the oncogenic activities *in vivo*, we performed the subcutaneous tumorigenesis assay and the metastasis assay by orthotopic transplantation. As a result, NMuMG-HOXB7 cells formed tumors when subcutaneously injected into immunodeficient mice (Figure 4a). Tumor formation was also observed in orthotopic transplantation. Primary tumors at fatpad were formed in half of mice within one week and in almost all mice within ten days (Figure 4b). They continued to grow until sixteen weeks after transplantation (Figure 4c), suggesting that *HOXB7* induced oncogenic activities *in vivo*. Next, we examine the metastatic activity of NMuMG-luc-HOXB7 cells that express luciferase gene with *HOXB7* gene. As a result of *in vivo* bioluminescence imaging with higher sensitivity, bioluminescence from the lung was detected in one mouse of twelve at sixteen weeks after transplantation (Figure 4d). To detect lung metastasis with more sensitivity, we excised the lungs and performed *ex vivo* bioluminescence imaging. As a result, lung metastasis was detected in 4 of 12 mice (Figure 4e). Our results showed that *HOXB7* induced not only *in vivo* tumor formation but also lung metastasis in orthotropic transplantation model.

**Figure 4.**
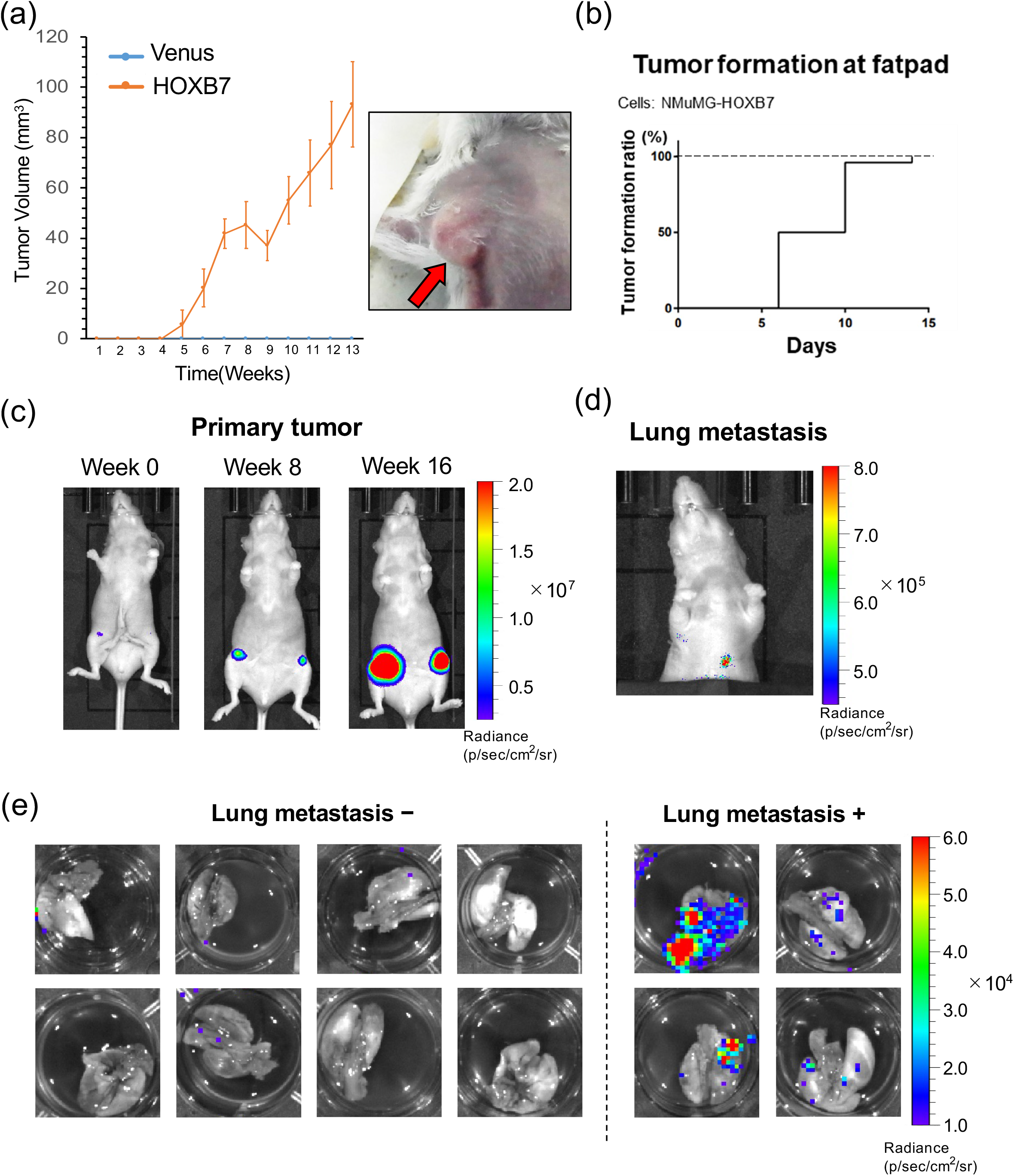
Overexpression of *HOXB7* induced tumorigenesis and lung metastasis. (a) Tumorigenesis assay of *HOXB7*-expressing NMuMG cells. *Venus*-expressing NMuMG cells were used as a negative control. Left: Tumor growth curve with volume. *HOXB7*-expressing NMuMG cells formed tumors following subcutaneously transplantation of *Rag2*^-/-^ mice (n = 10). Right: Image of *HOXB7*-positve tumor. (b) Primary tumor formation curve by orthotopic transplantation. *HOXB7*-expressiong NMuMG-luc cells were transplanted to BALB/cSlc-nu/nu mice (n = 12). (c) Representative *in vivo* luminescence images of primary tumor in orthotopic transplantation. (d) Representative *in vivo* luminescence image of lung metastasis in orthotopic transplantation. (e) *Ex vivo* luminescence images of lungs excised from orthotopically transplanted mice. Left: Lungs with no lung metastasis. Right: Lungs with lung metastasis.

To examine the contribution of STAT3 to *HOXB7*-mediated tumorigenicity *in vivo*, we performed the knock down (KD) of STAT3 in NMuMG-luc-HOXB7 cells. STAT3 KD cells were established using shRNA with retroviral vector (Figure 5a). As a result, STAT3 KD using shSTAT3#3 suppressed their focus-forming activity (Figure 5b). Next, STAT3 KD using shSTAT3#3 significantly decreased tumorigenesis at orthotopic sites (Figures 5c and 5d). Our results indicated that STAT3 signaling contributed to tumorigenesis induced by *HOXB7*.

**Figure 5.**
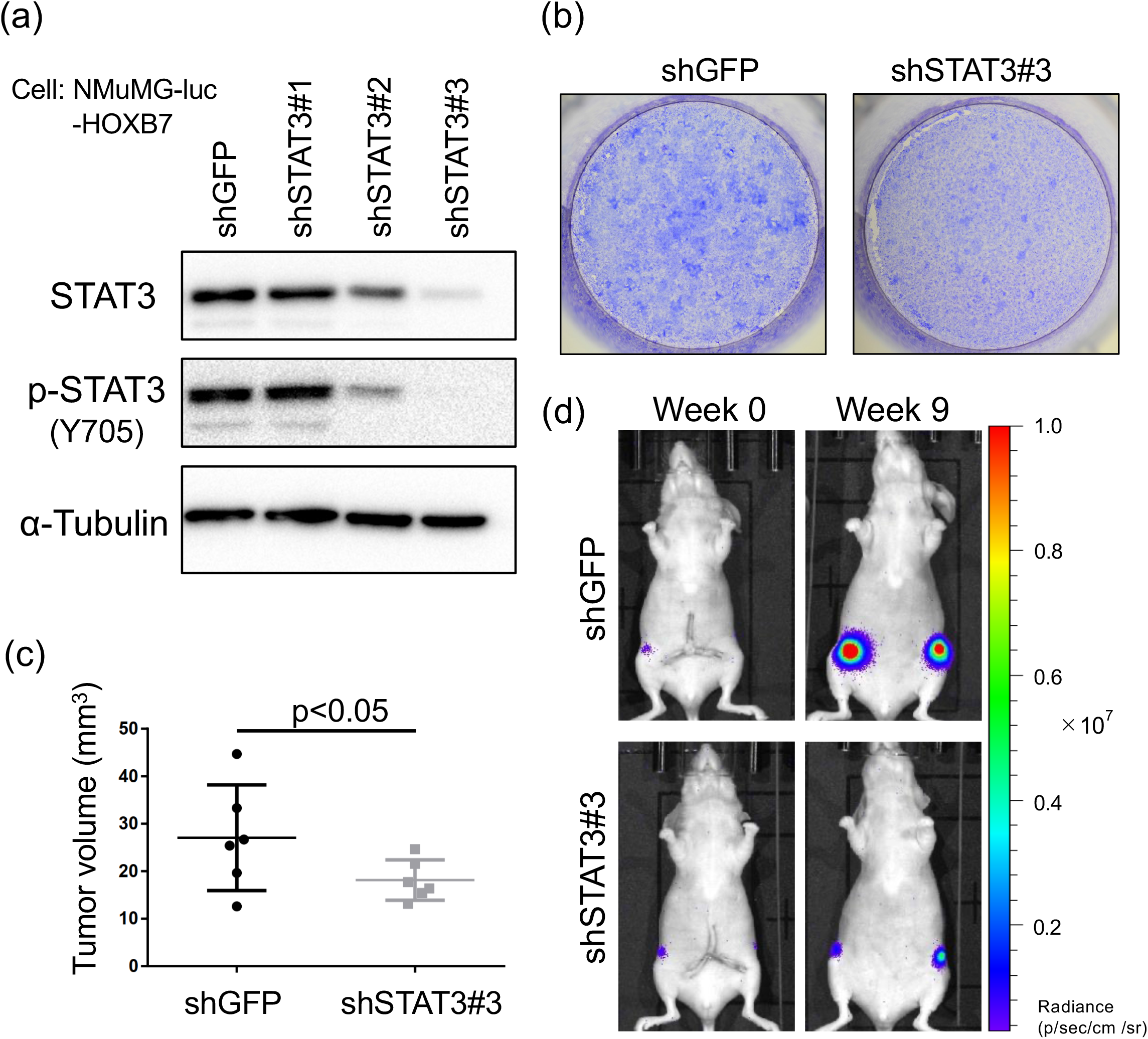
STAT3 contributed to tumorigenicity of NMuMG-HOXB7 cells. (a) Western blotting of STAT3 expression with shSTAT3 knockdown. (b) Focus formation assay of STAT3 knock down in NMuMG-HOXB7 cells. Foci were stained with crystal violet. Photos were taken at 10 days after the start of the experiment (n = 3 biological replicates; 2 independent experiments). (c) Tumor volume of primary tumor in orthotopic transplantation using NMuMG-luc-HOXB7-shGFP cells or NMuMG-luc-HOXB7-shSTAT3#3 cells. Student’s t-test (p < 0.05, n = 6). (d) Representative *in vivo* luminescence images of primary tumor in orthotopic transplantation using NMuMG-luc-HOXB7-shGFP cells or NMuMG-luc-HOXB7-shSTAT3#3 cells to BALB/cSlc-nu/nu mice (n = 3).

## 4. Discussion

In this study, we demonstrated that *HOXB7* induced cellular transformation and tumorigenesis to metastasis in transplantation models through activation of the JAK-STAT pathway. These results together with coamplification of *HOXB7* and *ERBB2* support its role in the poor prognosis of HER2-positive breast cancer patients.

HOX family genes regulate the development and homeostasis as transcriptional factors (Bhatlekar *et al*. 2014; Rezsohazy *et al*. 2015; Li *et al*. 2019). Several HOX genes contribute to cancer malignancy. In fact, overexpression or knockdown experiments supported the idea of *HOXB7* as an oncogene in the cases of breast, prostate, hepatocellular and gastric cancer cell lines (Errico *et al*. 2016; He *et al*. 2017; Wang *et al*. 2017). *HOXB7* also induced EMT and migration/invasion activity in MCF10A normal human mammary epithelial cells (Wu *et al*. 2006). *In vivo, HOXB7* promoted malignant progression through activation of the TGFß signaling in a MMTV-Hoxb7/Her2 transgenic mouse model (Liu *et al*. 2015). In addition, *HOXB7* overexpression in MCF7 cells (luminal A subtype) promoted acquired resistance to tamoxifen via activation of receptor tyrosine kinases and ERα signaling (Jin *et al*. 2012; Jin *et al*. 2015). Previous analysis of microarray data showed that high levels of *HOXB7* was associated with poor prognosis in HER2–positive, but not in HER2–negative breast cancers (Chen *et al*. 2008). Consistently, our analysis with recent genomic and RNA-seq data confirmed their study and further demonstrated that *HOXB7* was significantly co-amplified with *ERBB2*. Taken together, coamplification of *HOXB7* with *ERBB2* may enhance the malignancy of HER2-positive breast cancer.

Using an immortalized mammary gland epithelial cell NMuMG, *HOXB7* induced cellular transformation *in vitro* and tumor formation *in vivo*. In this cellular context, *HOXB7* increased the mRNAs of *Stat3* and protein levels of Jak2 and Stat3 with increased level of phosphorylated Stat3 and Stat5. Intriguingly, treatment with ruxolitinib, a JAK1/2 inhibitor, abolished cellular transformation with inhibition of Stat3 and Stat5 phosphorylation, suggesting the involvement of the JAK-STAT pathway in *HOXB7*-mediated transformation. Ruxolitinib binds intermediated-active kinase conformation of Jak2 (Kesarwani *et al*. 2015). As observed in previous research, it causes upregulation of Jak2 phosphorylation by ruxolitinib treatment via SOCS system (Lee *et al*. 2018). This phenomenon was also observed in the treatment with pacritinib, another JAK2 inhibitor (Hart *et al*. 2011). The JAK-STAT pathway is activated in several cancers and is believed to be involved in tumorigenesis (Hu *et al*. 2021). For this reason, kinase inhibitors against JAK including ruxolitinib have been developed. Even though *HOXB7* itself cannot be targeted, a crucial downstream JAK-STAT pathway may be targeted against cancer with *HOXB7* aberrations (Qureshy *et al*. 2020). In addition, since *Jak2* mRNA was not changed by *HOXB7* expression (Figure 2d), Jak2 is thought to be regulated at posttranscriptional level. The further analysis for expressional mechanisms of JAK-STAT signaling is needed for understanding the tumorigenesis induced by *HOXB7*.

Previous study demonstrated that *HOXB7* was not enough to induce tumor in *MMTV-Hoxb7* transgenic mice (Chen *et al*. 2008). In contrast, we showed that *HOXB7* alone was enough to induce tumorigenesis and lung metastasis in the context of NMuMG cells. This discrepancy cannot be explained, however, immortalization status of NMuMG may be involved. Nevertheless, our observation that *HOXB7* enhances lung metastasis in orthotropic model underpin the importance of *HOXB7* in lung metastasis, consistently with previous study of M*MTV-Hoxb7/Erbb2* transgenic mice. The transplantation models using NMuMG-luc-HOXB7 cells will be useful to discover the role of HOXB7 in cancer initiation and metastasis (Nakayama *et al*. 2021). Combined therapy against both *HOXB7* downstream JAK-STAT and ERBB2 may lead to efficacious outcome in “*ERBB2-* and *HOXB7*-amplified” subtype of breast cancer.

## Abbreviations

HOXB7: homeobox B7
NMuMG: normal murine mammary gland
HER2: human epidermal growth factor receptor 2
TCGA: the cancer genome atlas
TGFß: transforming growth factor beta
EMT: epithelial to mesenchymal transition
DMEM: Dulbecco’s modified Eagle’s medium
FBS: fetal bovine serum
IRES: internal ribosome entry site
PEI: polyethylenimine
METABRIC: the molecular taxonomy of breast cancer international consortium
IVIS: in vivo imaging system
SOCS: Suppressor of cytokine signaling.

## Acknowledgments

We thank Dr. Y. Yamamoto (National Cancer Center Research Institute) and Dr. J. Fujimoto (JBic) for the meaningful comments and discussion, Prof. K. Miyazawa (University of Yamanashi, Yamanashi, Japan) for kindly providing the NMuMG cell line, and Prof. T. Kitamura (Institute of Medical Science, The University of Tokyo, Tokyo, Japan) for kindly providing the Plat-E cell line. This study was conducted in part as a program through the Fukushima Translational Research Project.

## Funding

This study was supported by JSPS KAKENHI (grant no. 23241064, Grant-in-Aid for Scientific Research (A) to K.S.; grant no. 18K16269: Grant-in-Aid for Early-Career Scientist to J.N.; grant no. 20J01794, Grant in Aid for JSPS fellows to J.N.) and the Japan Biological Informatics Consortium (JBic).

## Authors’ contributions

J.N. and K.S. conceived and designed the study. K.A, M.S., S.K., A.M., and J.N. performed the experiments. K.A., M.S., K.N. and J.N. analyzed the data. N.G., S.W. and K.S. interpreted the data. J.N. wrote the manuscript. All authors reviewed and edited the manuscript.

## Conflict of interest

The authors declare no conflict of interest.

## Notes

### Competing Interest Statement

The authors have declared no competing interest.

### Summary of Updates

In the revised manuscript, we performed the knock down of STAT3 of NMuMG-HOXB7 cells.

