## Supplementray Figures and Tables for "*HOXB7* induces STAT3-mediated transformation and lung metastasis in immortalized mammary gland NMuMG cells"

### Supplementary Table 1. Antibody information

| Antibody | Maker | Catalog number | dilution | Blocking buffer |
|---|---|---|---|---|
| Phospho-Stat3 (Tyr705) (D3A7) XP® Rabbit mAb | CST | 9145 | 1:1000 | 5% BSA / 0.5% TBS-Tween |
| Stat3 (124H6) Mouse mAb | CST | 9139 | 1:1000 | 5% BSA / 0.5% TBS-Tween |
| Phospho-Jak1 (Tyr1034/1035) Antibody | CST | 3331 | 1:1000 | 5% BSA / 0.5% TBS-Tween |
| Jak1 (6G4) Rabbit mAb | CST | 3344 | 1:1000 | 5% BSA / 0.5% TBS-Tween |
| Phospho-Jak2 (Tyr1007/1008) Antibody | CST | 3771 | 1:1000 | 5% BSA / 0.5% TBS-Tween |
| Jak2 (D2E12) XP® Rabbit mAb | CST | 3230 | 1:1000 | 5% BSA / 0.5% TBS-Tween |
| Phospho-p44/42 MAPK (Erk1/2) (Thr202/Tyr204) (D13.14.4E) XP® Rabbit mAb | CST | 4370 | 1:2000 | 5% BSA / 0.5% TBS-Tween |
| p44/42 MAPK (Erk1/2) (137F5) Rabbit mAb | CST | 4695 | 1:2000 | 5% BSA / 0.5% TBS-Tween |
| Phospho-Akt(Thr)(C31E5E) Rabbit mAb | CST | 2965 | 1:2000 | 5% BSA / 0.5% TBS-Tween |
| Phospho-Akt(Ser473)(D9E)XP Rabbit mAb | CST | 4060 | 1:2000 | 5% BSA / 0.5% TBS-Tween |
| Akt (pan) (C67E7) Rabbit mAb | CST | 4691 | 1:2000 | 5% BSA / 0.5% TBS-Tween |
| Stat1 Antibody | CST | 9172 | 1:1000 | 5% BSA / 0.5% TBS-Tween |
| Phospho-Stat1 (Ser727) Antibody | CST | 9177 | 1:1000 | 5% BSA / 0.5% TBS-Tween |
| Stat5 Antibody | CST | 9363 | 1:1000 | 5% BSA / 0.5% TBS-Tween |
| Phospho-Stat5 (Tyr694) (D47E7) XP® Rabbit mAb | CST | 4322 | 1:1000 | 5% BSA / 0.5% TBS-Tween |
| Purified Mouse Anti-E-Cadherin | BD transduction | 610182 | 1:2000 | 5% skim milk /0.5% TBS-Tween |
| Purified Mouse Anti-N-Cadherin | BD transduction | 610921 | 1:2000 | 5% skim milk /0.5% TBS-Tween |
| Purified Mouse Anti-Vimentin | BD transduction | 550513 | 1:2000 | 5% skim milk /0.5% TBS-Tween |
| HoxB7 antibody (747C4a) | Santa Cruz | sc-81292 | 1:1000 | 5% skim milk /0.5% TBS-Tween |
| anti-α-Tubulin clone DM1A | Sigma | T6119 | 1:2000 | 5% skim milk /0.5% TBS-Tween |
| anti-mouse Ig, HRP-Linked Whole Ab Sheep | GE healthcare | NA931 | 1:5000 | 5% skim milk /0.5% TBS-Tween |
| anti-rabbit Ig, HRP-Linked Whole Ab Donkey | GE healthcare | NA934 | 1:5000 | 5% skim milk /0.5% TBS-Tween |

### Supplementary Table 2. Primers for qRT-PCR

| Target gene | Sequence 5'->3' |
|---|---|
| Stat3 Forward | GCACCTTGGATTGAGAGTCA |
| Stat3 Reverse | CCCAAGAGATTATGAAACACCA |
| Jak2 Forward | CATACACAGCAACTGCCACG |
| Jak2 Reverse | TTTTCTCGCTCAACAGCAAAGG |
| 18srRNA Forward | GTAACCCGTTGAACCCCATT |
| 18srRNA Reverse | CCATCCAATCGGTAGTAGCG |

Supplementary Figure 1

Raw data of western blotting in Figure 2C.

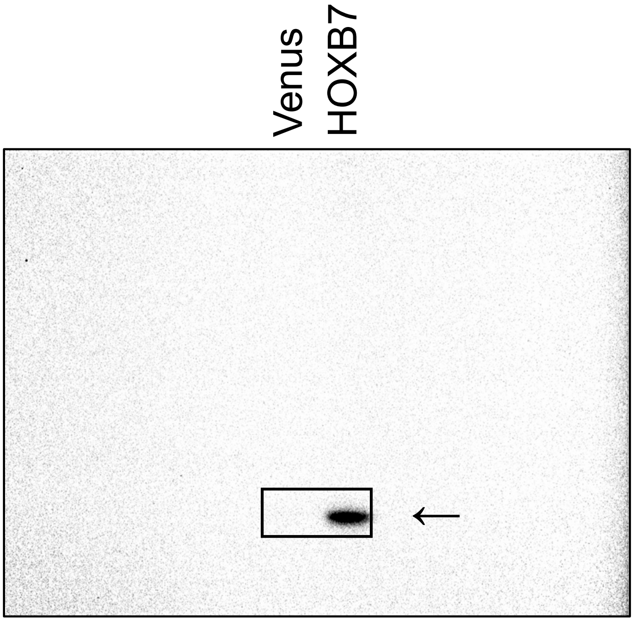


Fig2C anti-HOXB7

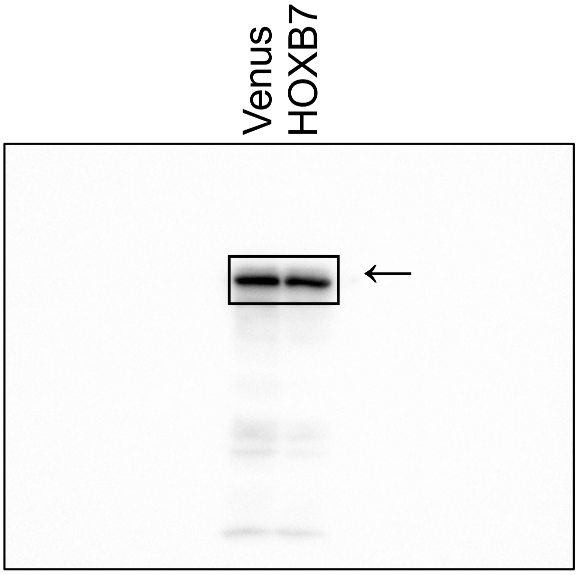


Fig2C anti-Ecadherin

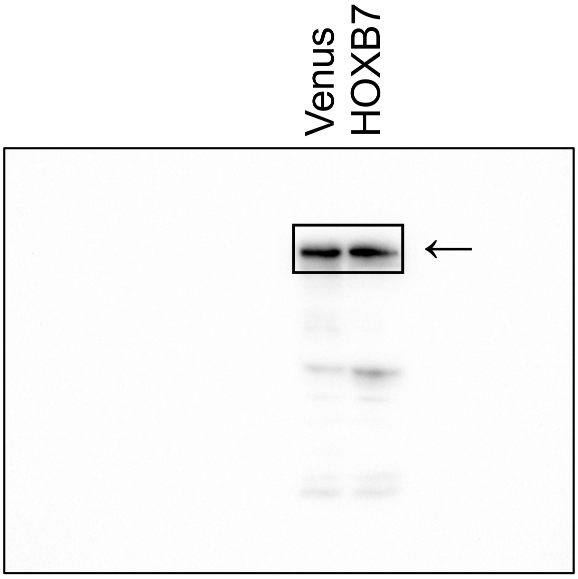


Fig2C anti-Ncadherin

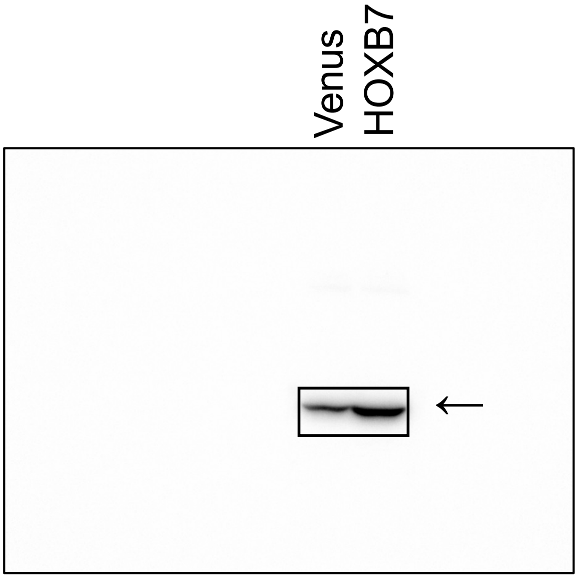


Fig2C anti-Vimentin

Supplementary Figure 1

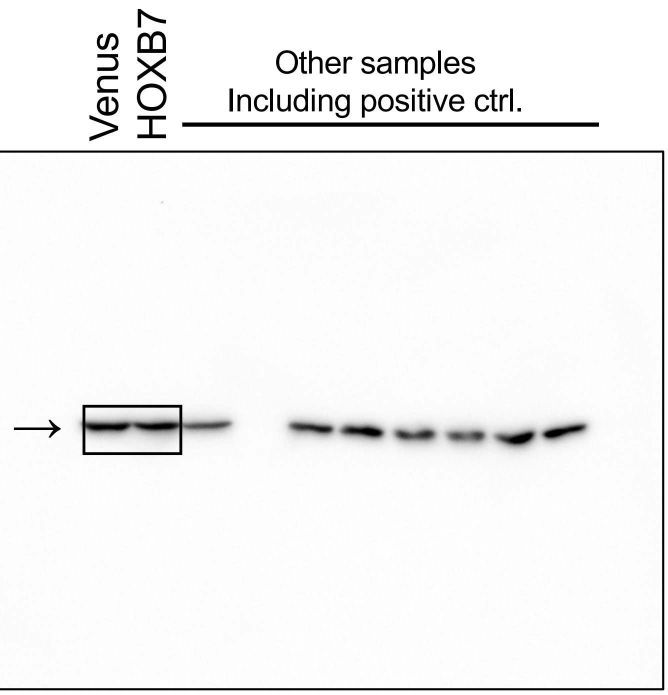


Fig2C anti-Tubulin

Supplementary Figure 1

Raw data of western blotting in Figure 2D.

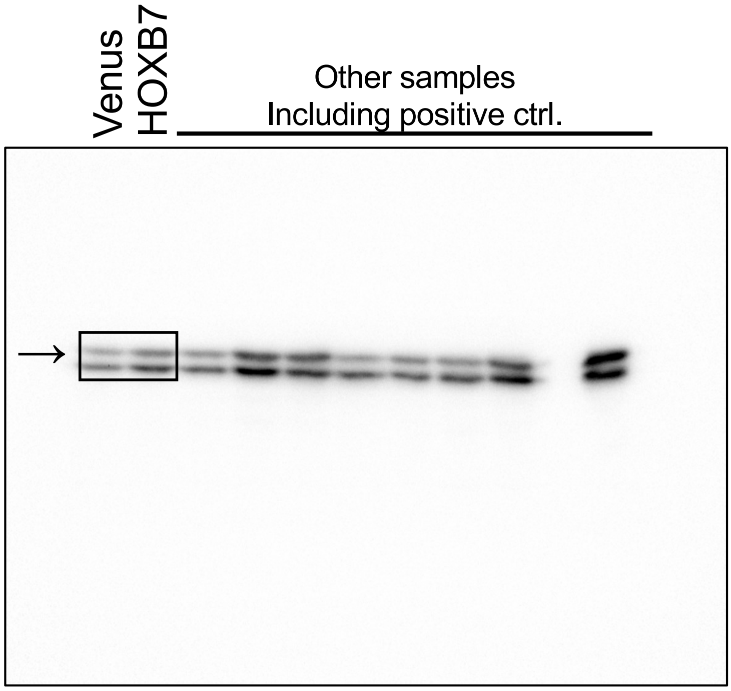


Fig2D anti-p-Erk

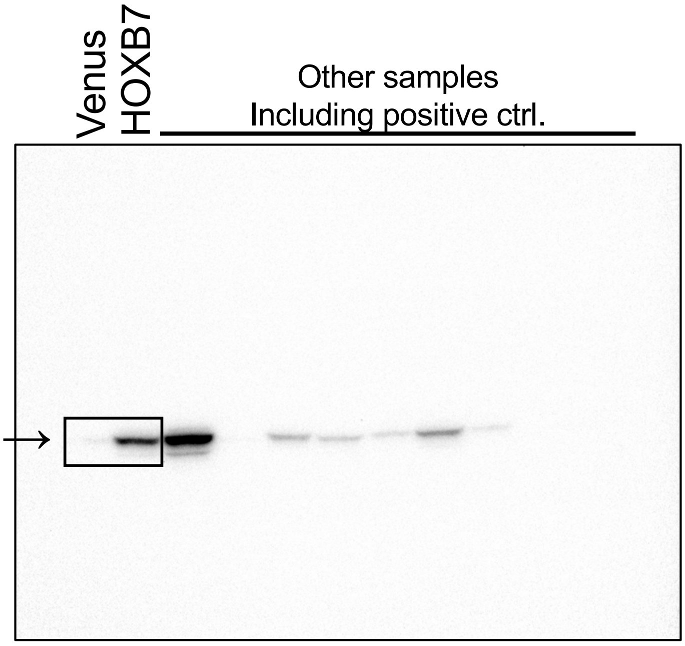


Fig2D anti-p-Stat3

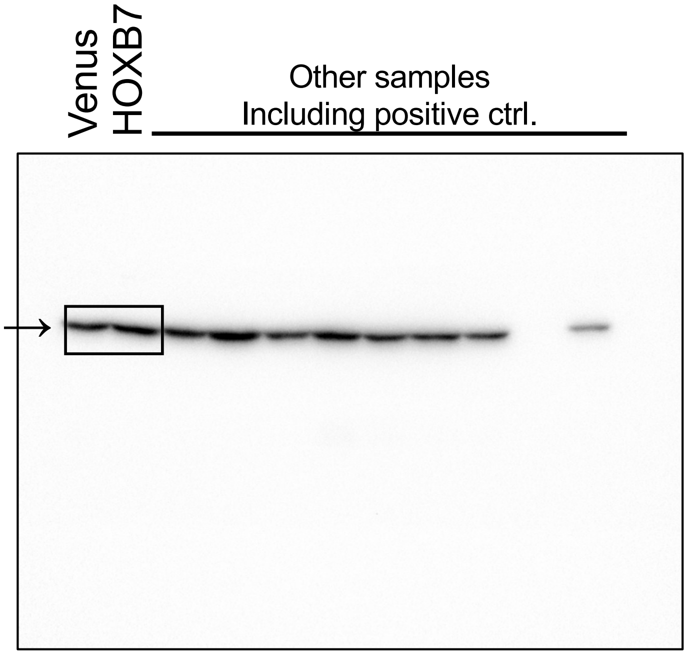


Fig2D anti-p-Akt(S473)

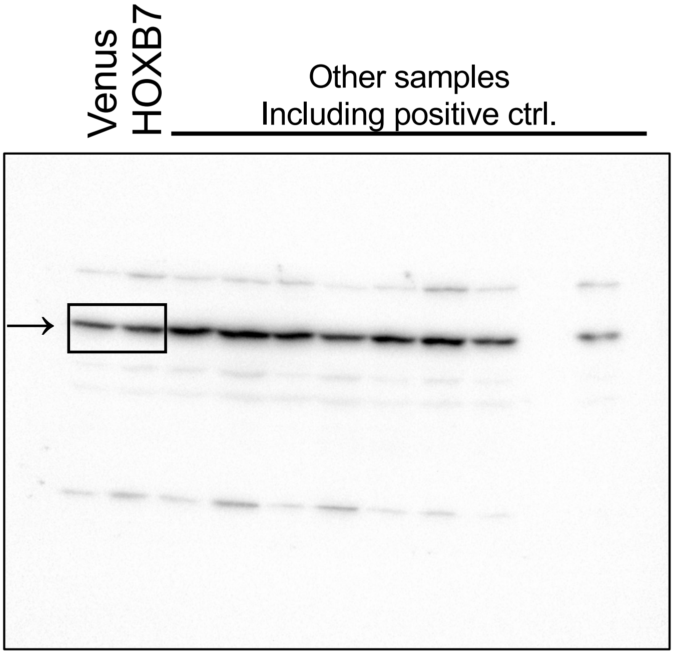


Fig2D anti-p-Akt(T308)

Supplementary Figure 1

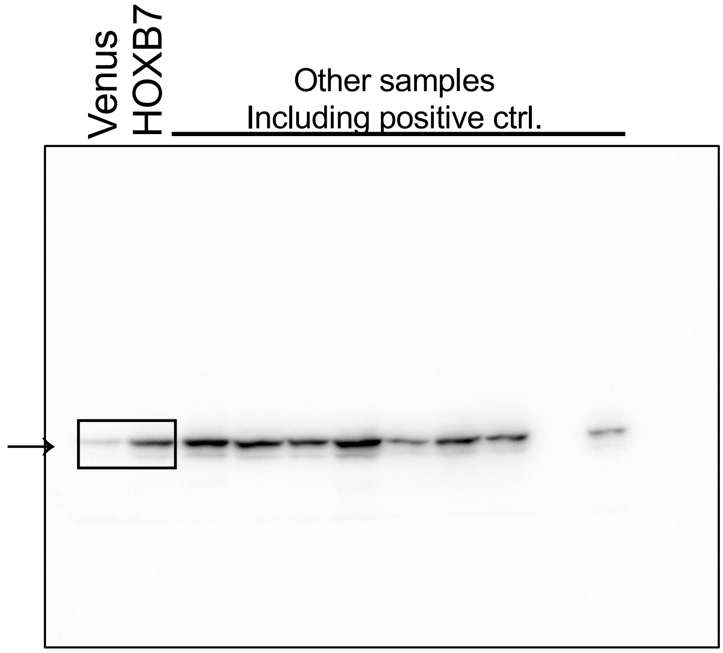


Fig2D anti-Stat3

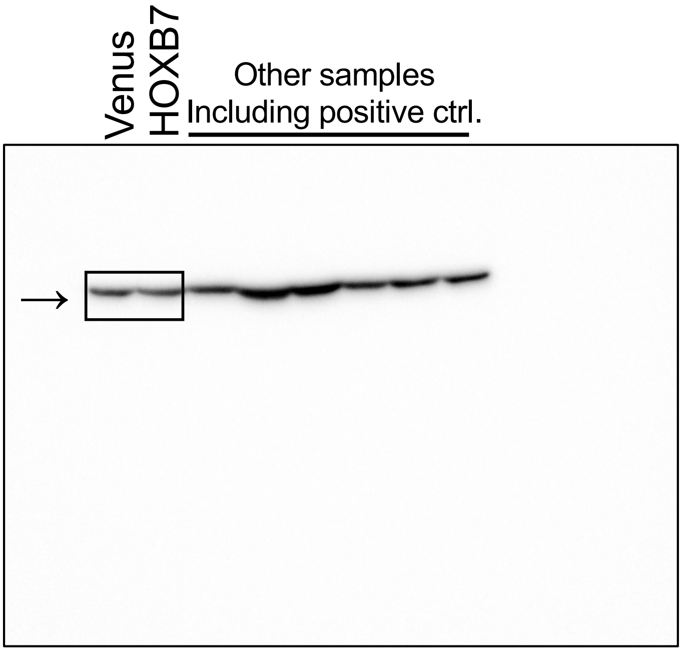


Fig2D anti-Tubulin

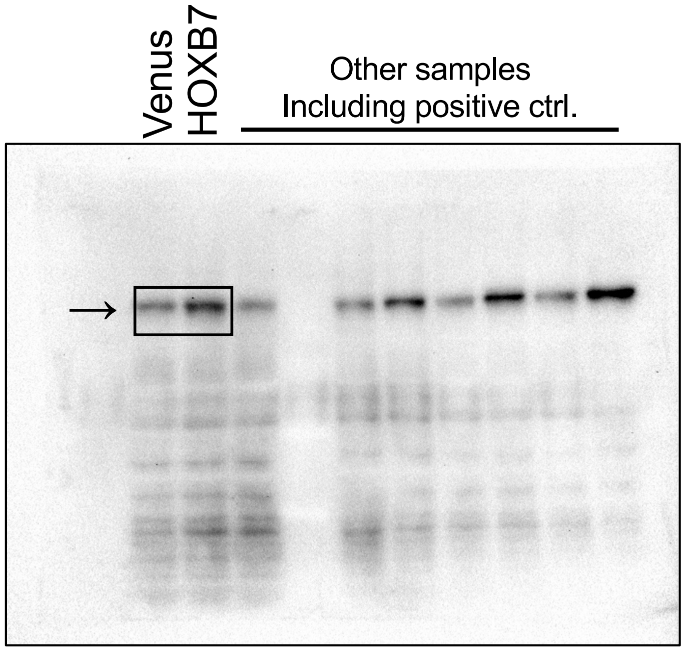


Fig2D anti-p-Jak2

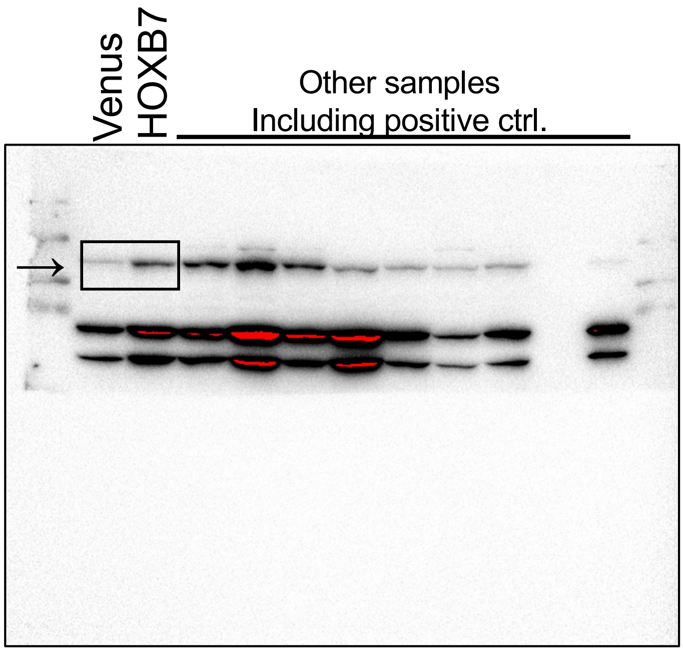


Fig2D anti-Jak2

Supplementary Figure 1

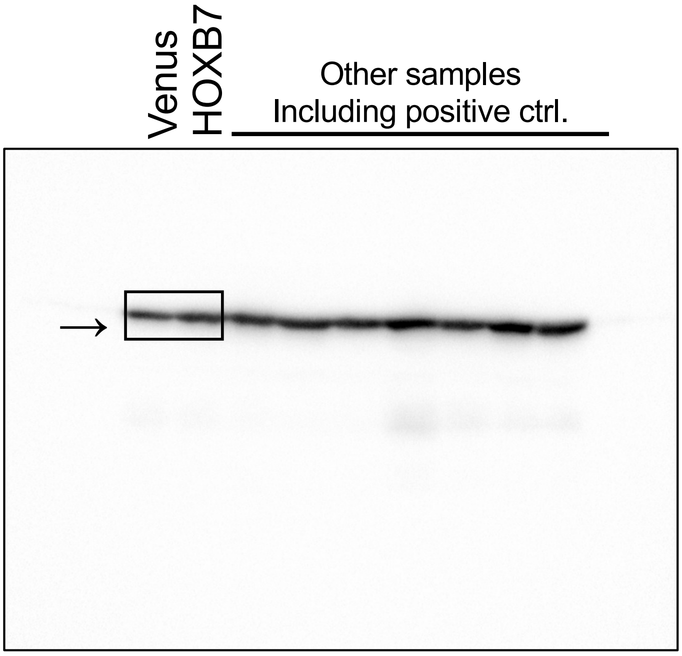


Fig2D anti-Akt

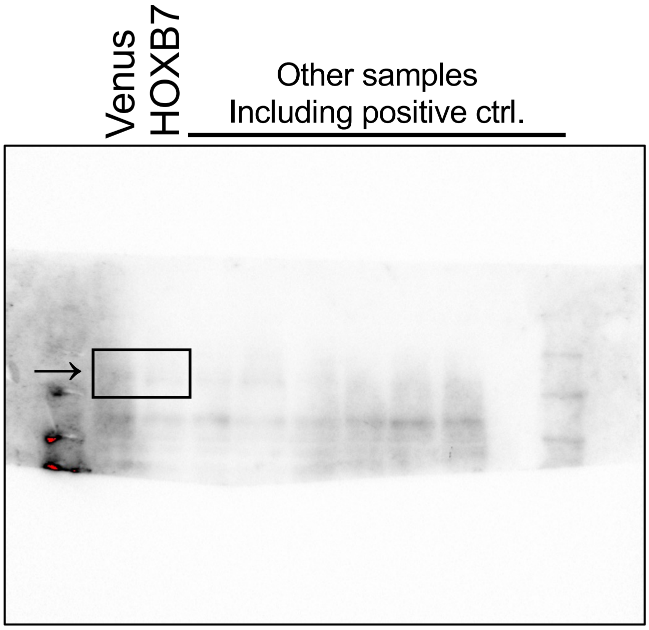


Fig2D anti-p-Jak1

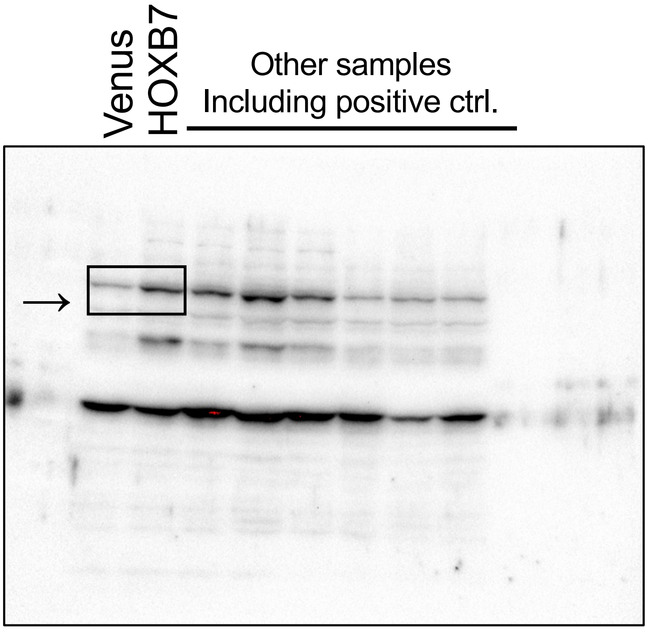


Fig2D anti-Jak1

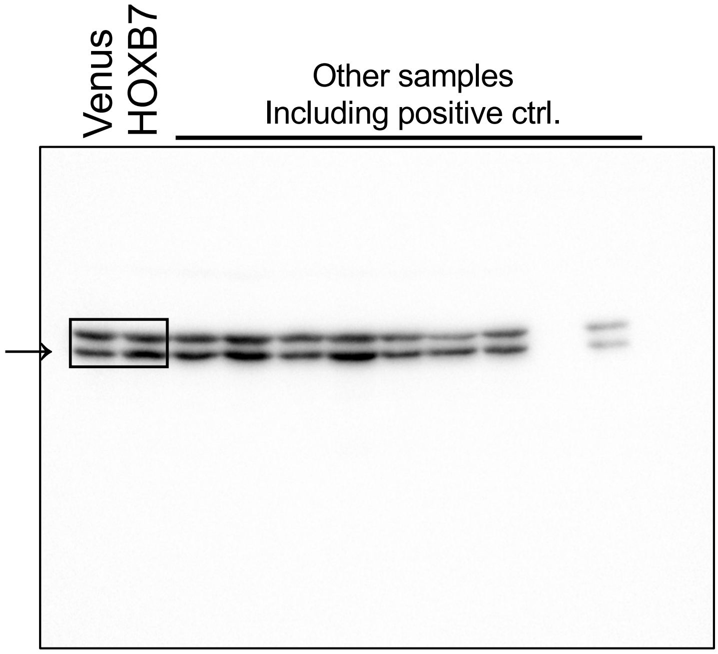


Fig2D anti-Erk

Supplementary Figure 1

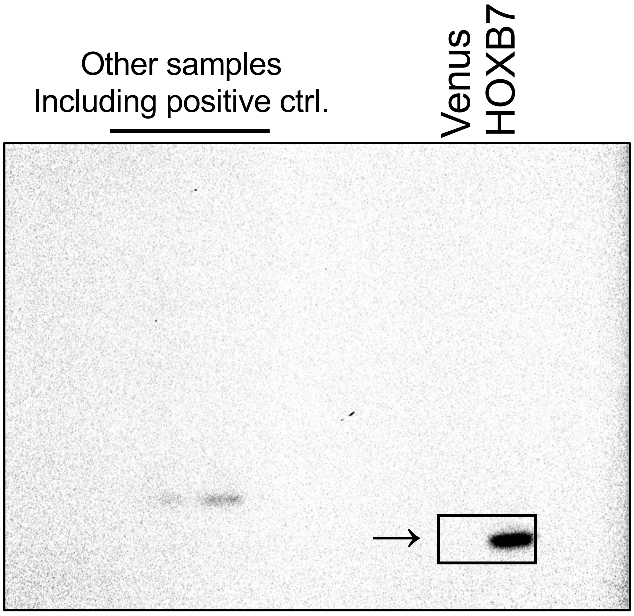


Fig2D anti-HOXB7

Supplementary Figure 1

Raw data of western blotting in Figure 3A.

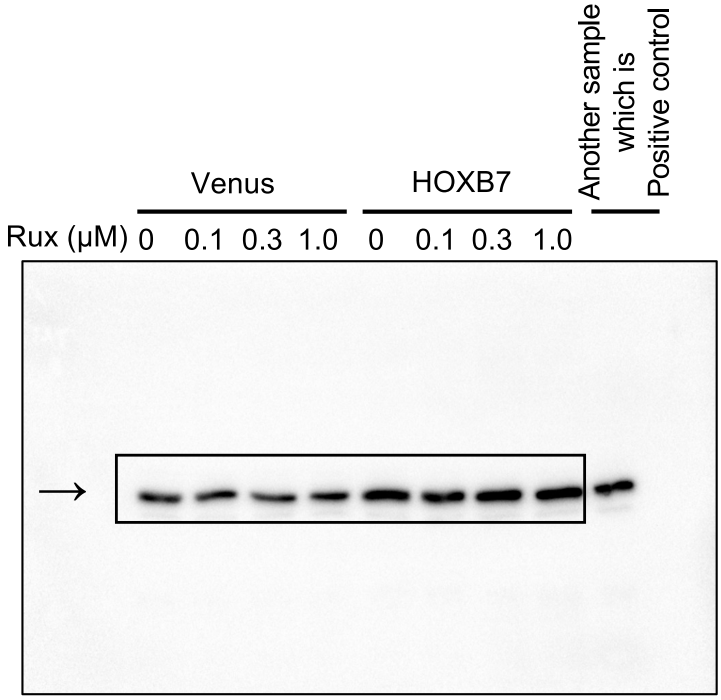


Fig3A anti-Stat3

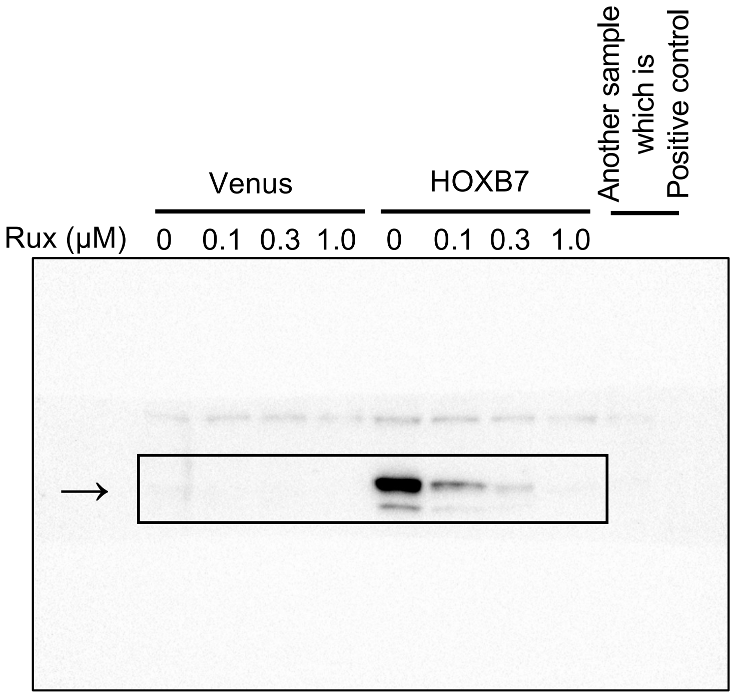


Fig3A anti-p-Stat3

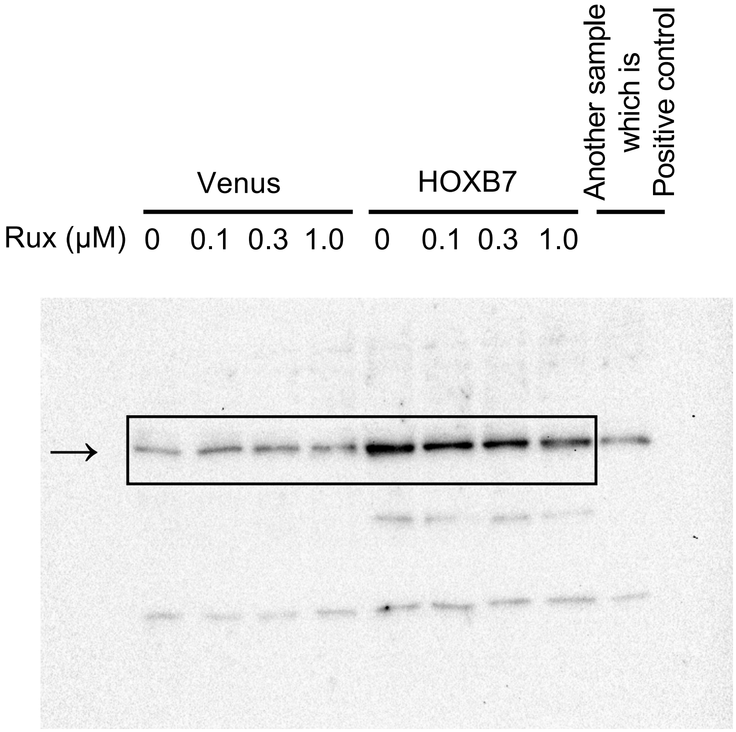


Fig3A anti-Jak1

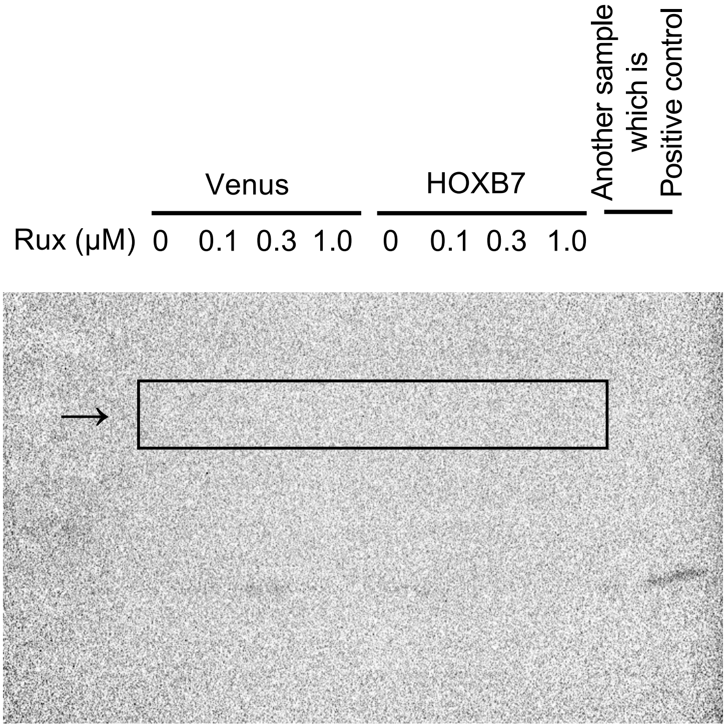


Fig3A anti-p-Jak1

Supplementary Figure 1

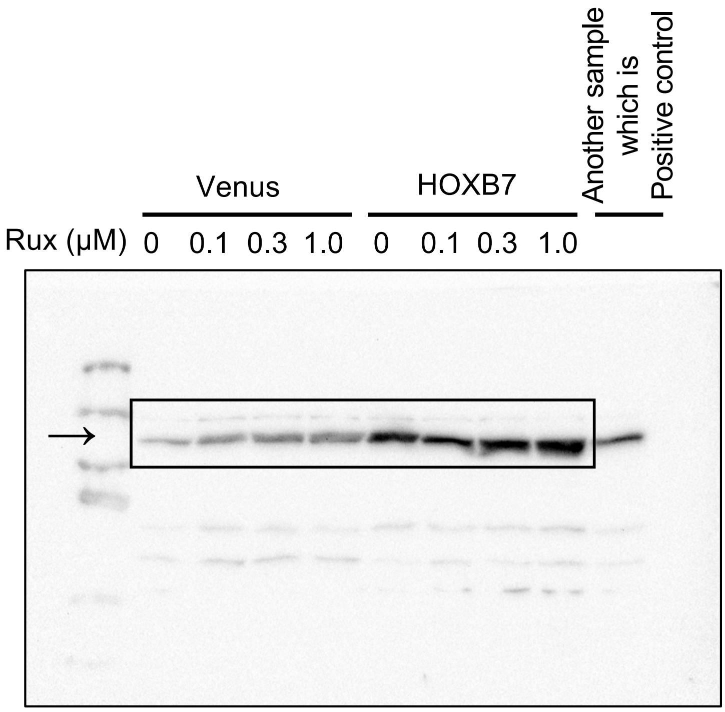


Fig3A anti-Jak2

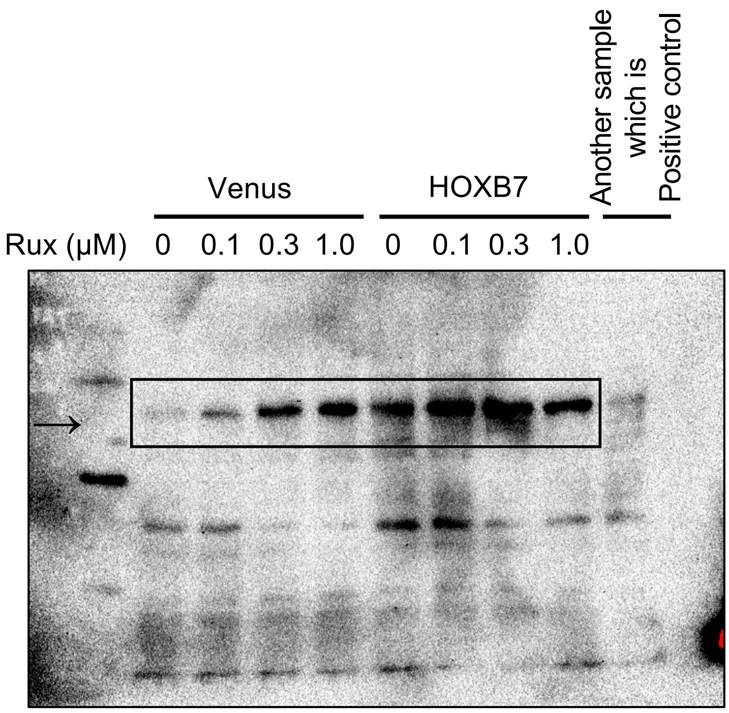


Fig3A anti-p-Jak2

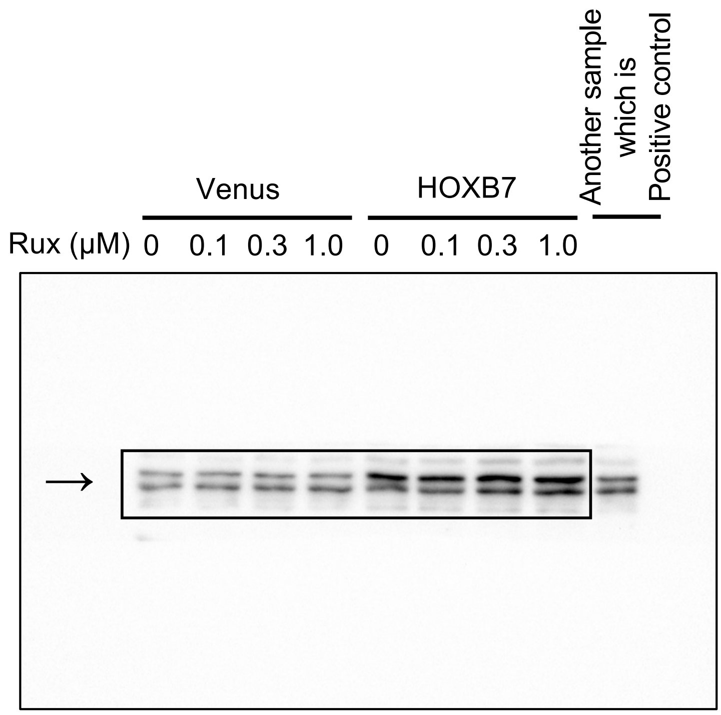


Fig3A anti-Stat5

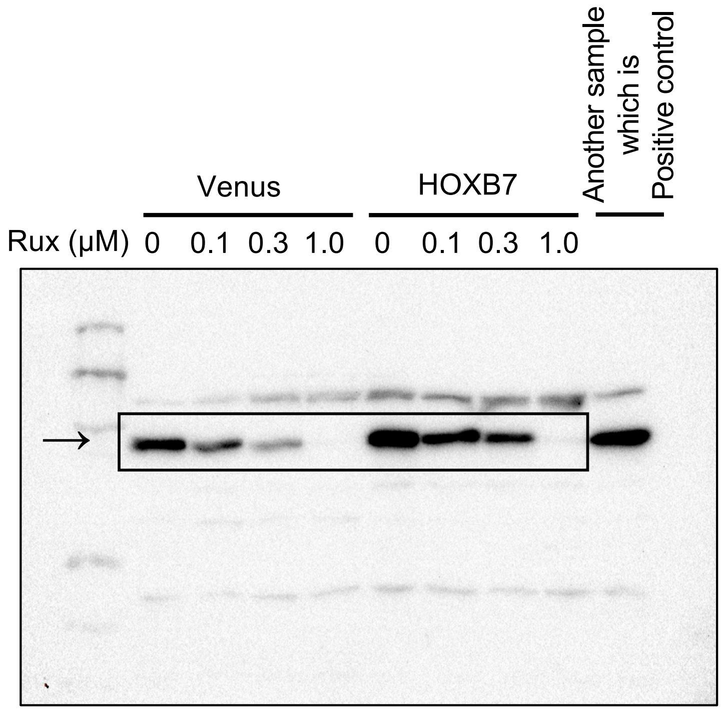


Fig3A anti-p-Stat5

Supplementary Figure 1

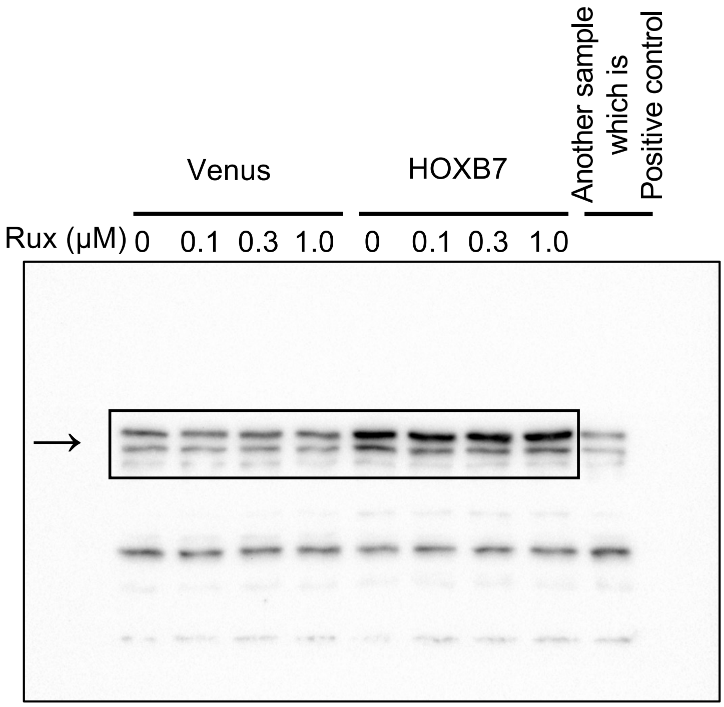


Fig3A anti-Stat1

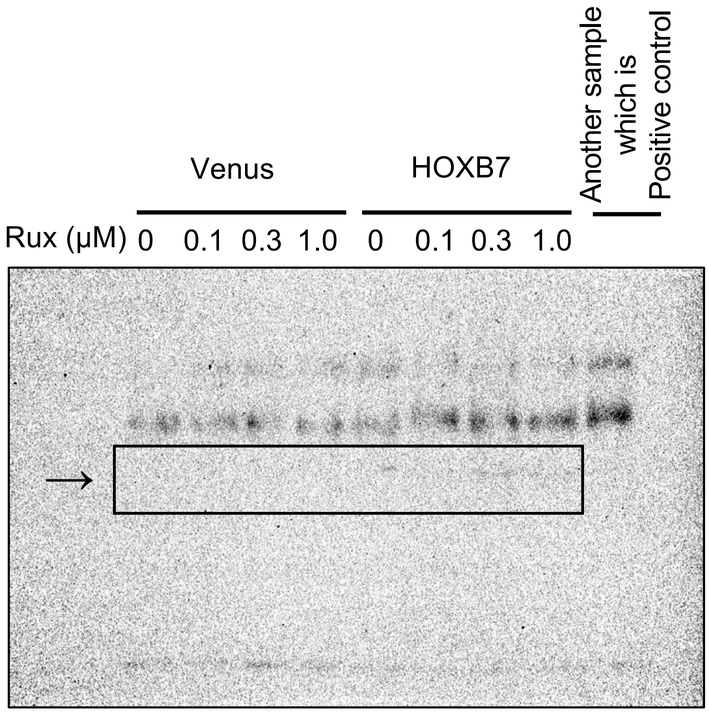


Fig3A anti-p-Stat1

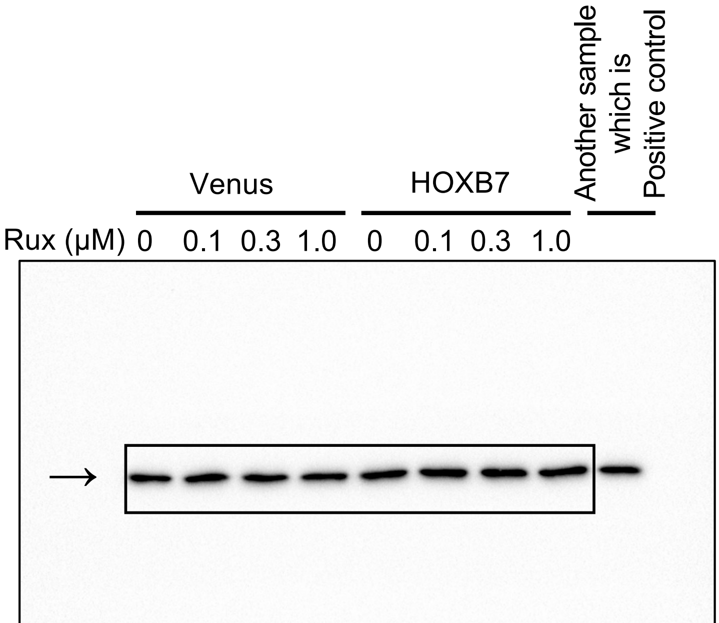


Fig3A anti-Tubulin

Supplementary Figure 1

Raw data of western blotting in Figure5A.

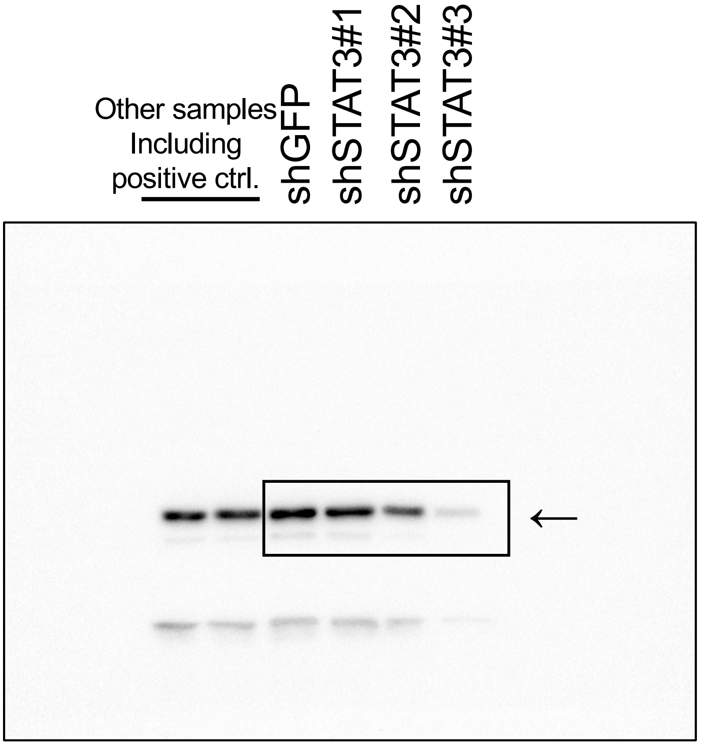


Fig5A anti-Stat3

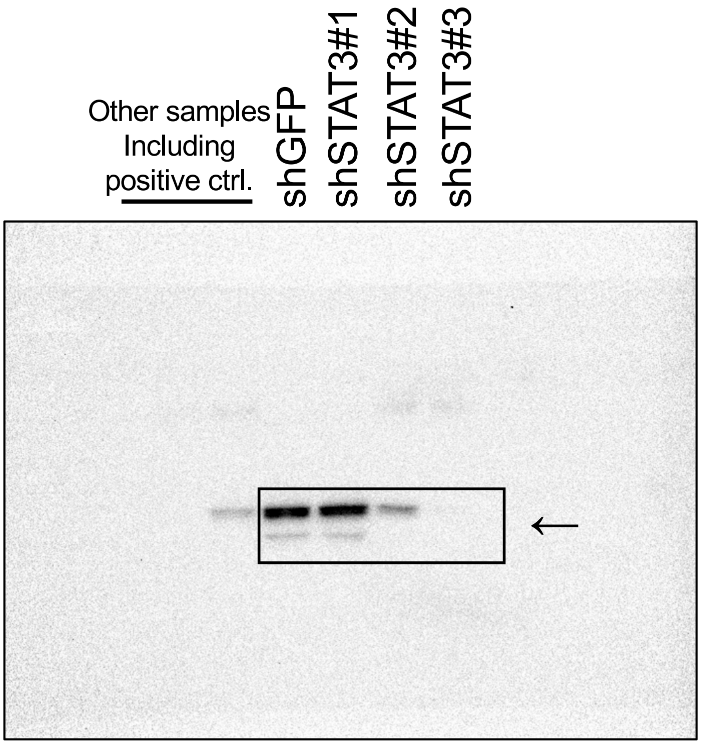


Fig5A anti-p-Stat3

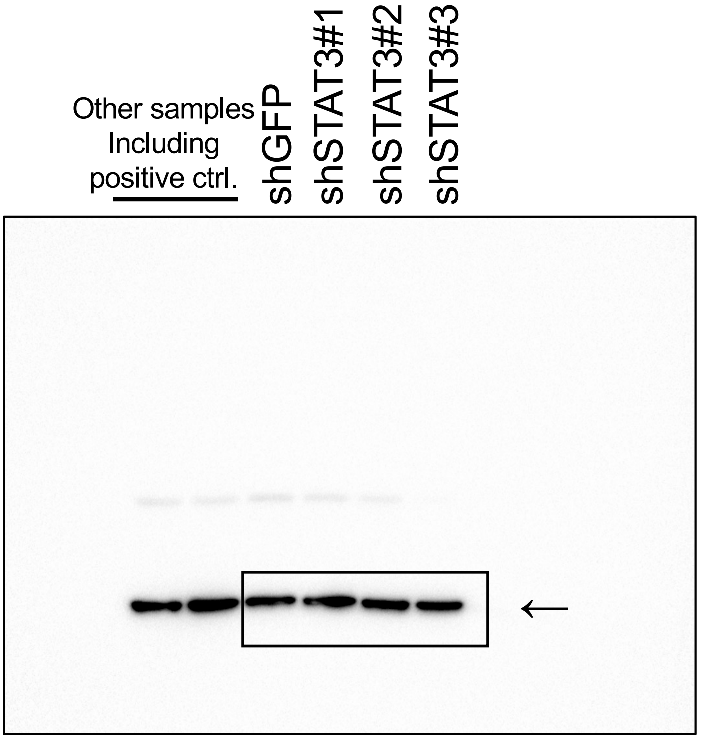


Fig5A anti-α-Tubulin

Supplementaly Fig2

Results of Focus formation assay in NMuMG-Luc-HOXB7-shSTAT3#2

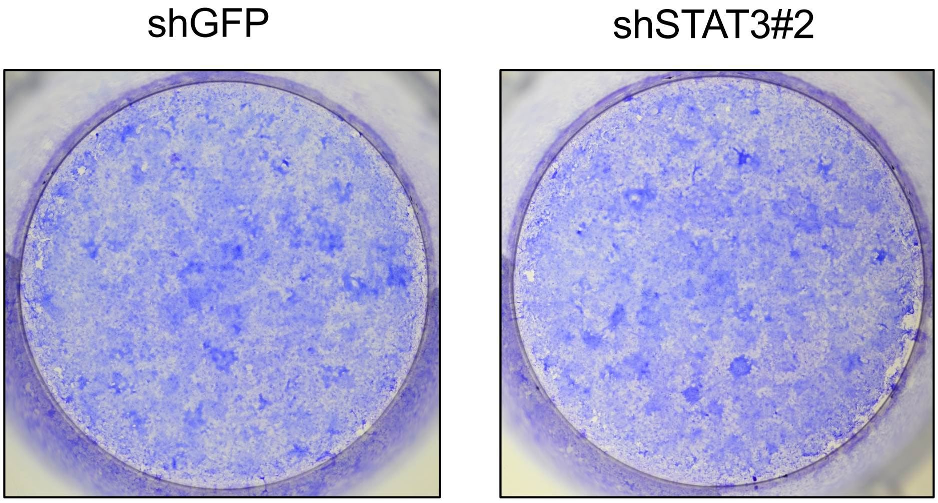

Supplementaly Fig2

Results of Orthotopic transplantation in NMuMG-Luc-HOXB7-shSTAT3#2

A

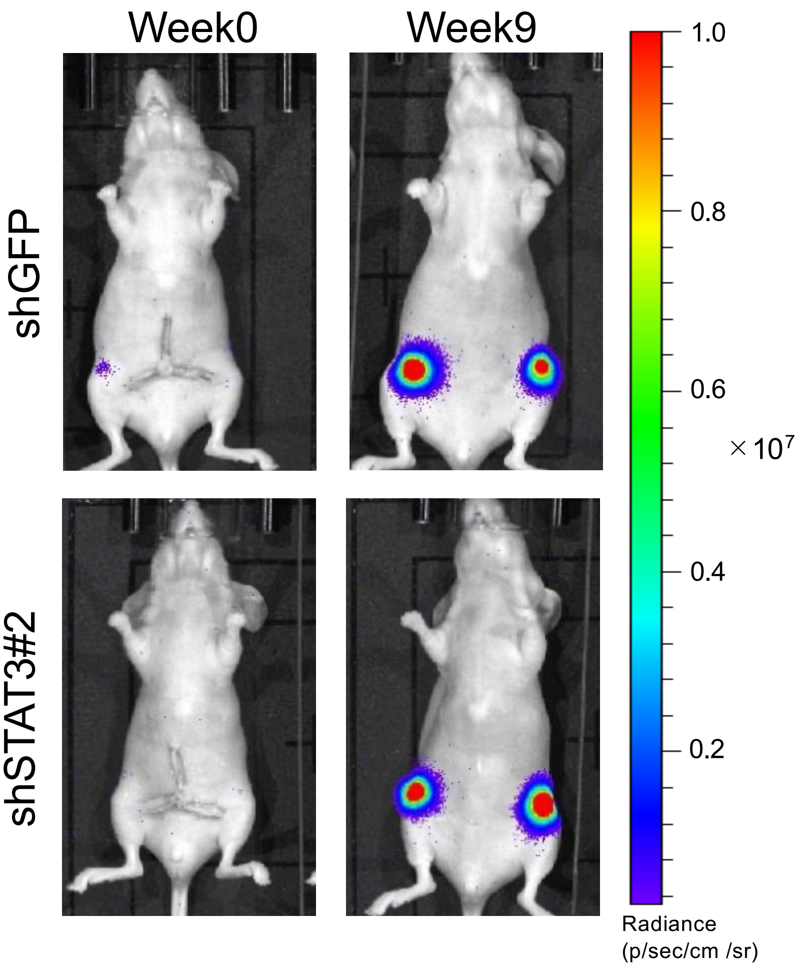


B

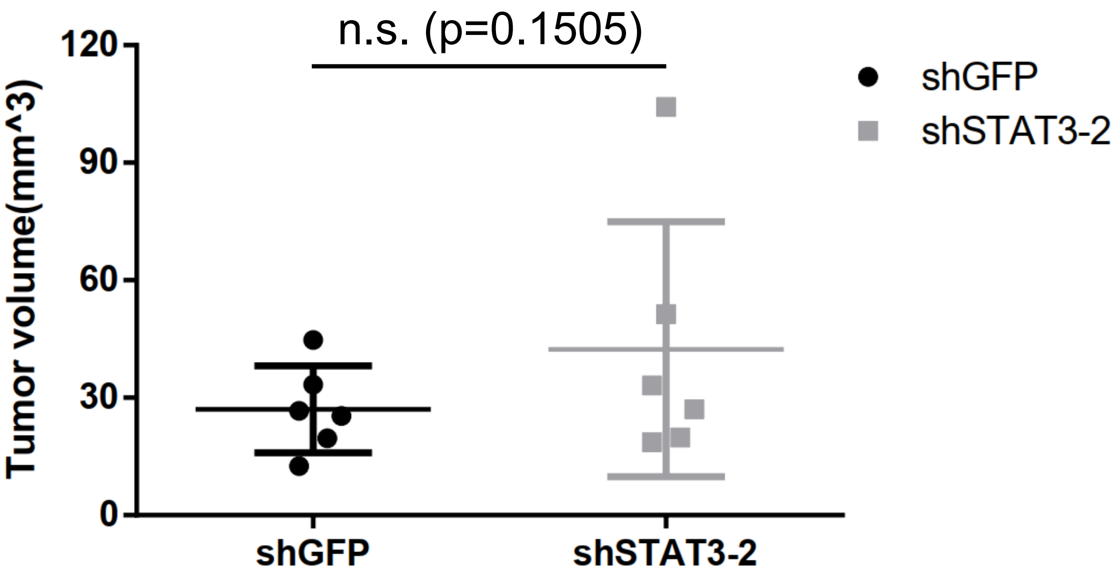
